# Increased microclimatic variation in artificial nests does not create ecological traps for a secondary cavity breeder

**DOI:** 10.1101/2020.05.14.095950

**Authors:** Timothée Schwartz, Arnaud Genouville, Aurélien Besnard

**Author notes:** Correspondence author: 51 rue des Palombes, 34670 St-Bres, France.

## Abstract

Artificial devices are increasingly used in conservation measures to mitigate the disappearance of natural habitats. However, few studies have demonstrated their benefits for the target species, and they may pose a risk of creating ecological traps. This occurs when lower individual fitness is found in artificial habitats that are more attractive than their natural equivalents. In this study, we tested the ecological trap hypothesis on a dense population of European rollers *Coracias garrulus* breeding in both natural cavities and nest-boxes. Our initial hypothesis was that the more stressful microclimatic conditions of nest-boxes would lead to reduced fitness of rollers, thus creating an ecological trap. The results showed that nest-boxes were preferred over natural cavities. Despite significantly more extreme microclimatic conditions in nest-boxes, we found similar breeding parameters between artificial and natural nest types. Our results also suggest that rollers selected the nest-boxes which best buffered the temperature, thus avoiding potential ecological traps. Overall our results lead to the conclusion that nest-boxes do not create ecological traps for rollers in this study site. However, other species may be more sensitive to microclimatic variations or less able to avoid the least favourable nest-boxes. These findings could help to inform the placement of nest-boxes in order to reduce extreme temperatures and variation in humidity rates. Future studies could compare nest types for other fitness parameters, such as juvenile body condition or survival. We also recommend the ecological trap hypothesis as a useful framework to evaluate the outcomes of artificial devices used for conservation.

## 1. Introduction

The rapid changes affecting biodiversity in recent decades have led to an expansion of habitat restoration and biodiversity offsetting programmes (Dobson, 1997; Maron, Gordon, Mackey, Possingham & Watson, 2015). These initiatives have resulted in the development of numerous methods to restore or recreate habitats that have been damaged or destroyed by human activities such as urbanization or intensive agriculture. Of these methods, the use of artificial devices for conservation is increasingly implemented as a response to habitat degradation. For example, shelters for reptiles are positioned to replace disappearing stone walls and hedges (Grillet et al., 2010), bat boxes to compensate for felled cavity trees or restored buildings (Flaquer, Torre & Ruiz-Jarillo, 2006), and nest-boxes to provide additional breeding places for birds and mammals (Goldingay & Stevens, 2009). Yet despite their extensive use in conservation and offsetting programmes, robust evaluations of the benefit of artificial devices in terms of the population viability of the target species are rare (Wesołowski, 2011, but see Bourgeois, Dromzée, & Vidal, 2015). The success of artificial devices is generally evaluated using single indicators, for example, colonization by the target species, (Aleman & Laurens, 2013; Avilés & Sanchez, 2000; Chapman & Blockley, 2009) or, in the best cases, their impact on some selected demographic parameters, such as breeding success or survival (see e.g. Bourgeois et al., 2015; Libois et al., 2012). However, measuring occupation alone does not necessarily demonstrate any benefit for the population. A fundamental issue is that while an artificial site may be attractive to the target species, it may have a deleterious impact on reproduction: for instance, by increasing predation risk (Robertson & Hutto, 2006). Furthermore, positive outcomes for the population are difficult to demonstrate unless demographic parameters (for instance, fecundity) in artificial devices are compared to those in natural breeding sites. Several studies comparing artificial devices alone in different contexts have shown that the fitness of the target species can decrease in some situations, such as inappropriate nest size (Demeyrier, Lambrechts, Perret, & Grégoire, 2016) or nest placement (Rodríguez-Ruiz, Avilés, & Parejo, 2011), hence creating an ecological trap (Klein, Nagy, Csorgo & Matics, 2007).

An ecological trap is a modified habitat that is preferred by a target species over an unmodified habitat, but which reduces the fitness of individuals (Robertson & Hutto, 2006). Ecological traps generally occur when perceived habitat quality of the modified habitat does not match its actual quality (Schlaepfer, Runge & Sherman, 2002). In the context of artificial devices, while some evaluations only explore either its attractiveness or its impact on the fitness of the target species, testing the ecological trap hypothesis requires to examine both (Robertson & Hutto, 2006). It also relies on comparing these parameters between the artificial device and the unmodified habitat that it aims to mimic. This latter aspect is, however, often lacking in evaluations of artificial conservation tools (but see Bourgeois et al., 2015). The test of the ecological trap hypothesis is thus a two-steps process (Fig. 2). First, studying a species’ preference for natural or artificial habitats is essential as avoidance of artificial devices would lead to the failure of a conservation strategy. Avoidance means that the target species is less attracted by the artificial habitat compared to its natural habitat and is generally the result of a lack of knowledge of the drivers of the target species’ habitat selection needed for relevant device conception or placement (Battin, 2004; Robertson & Hutto, 2006). The second step consists in measuring fitness parameters or proxies at the population level (for example, demographic parameters), which is often more challenging than for preference. These parameters must be indeed compared with known parameters for healthy populations of the same species living in an unmodified habitat in a similar context, however, such habitats are often difficult to find or access (Lambrechts et al., 2010).

**Figure 1:**
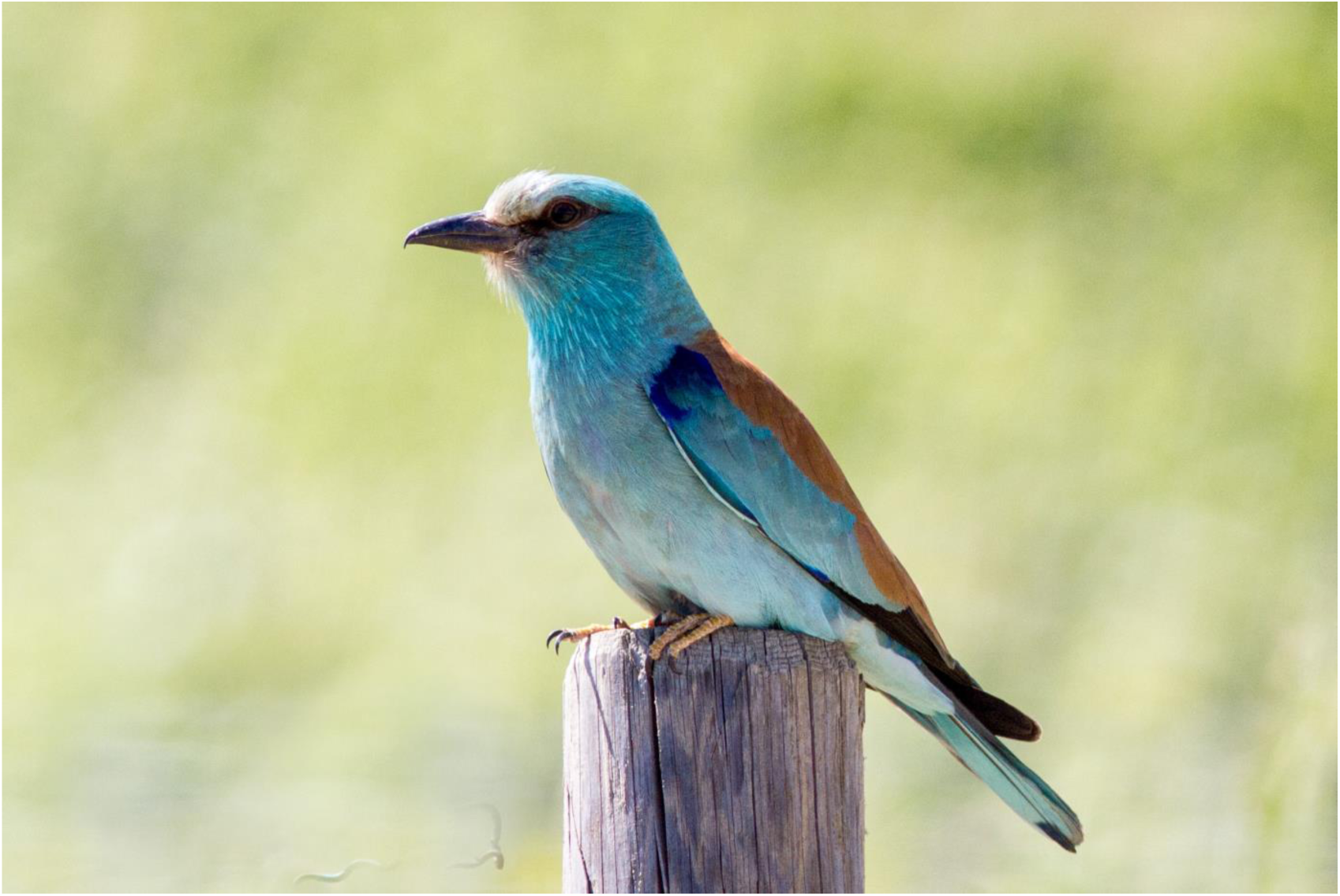
European roller *Coracias garrulus* in Vallée des Baux, France (Author: Peter Harris)

**Figure 2:**
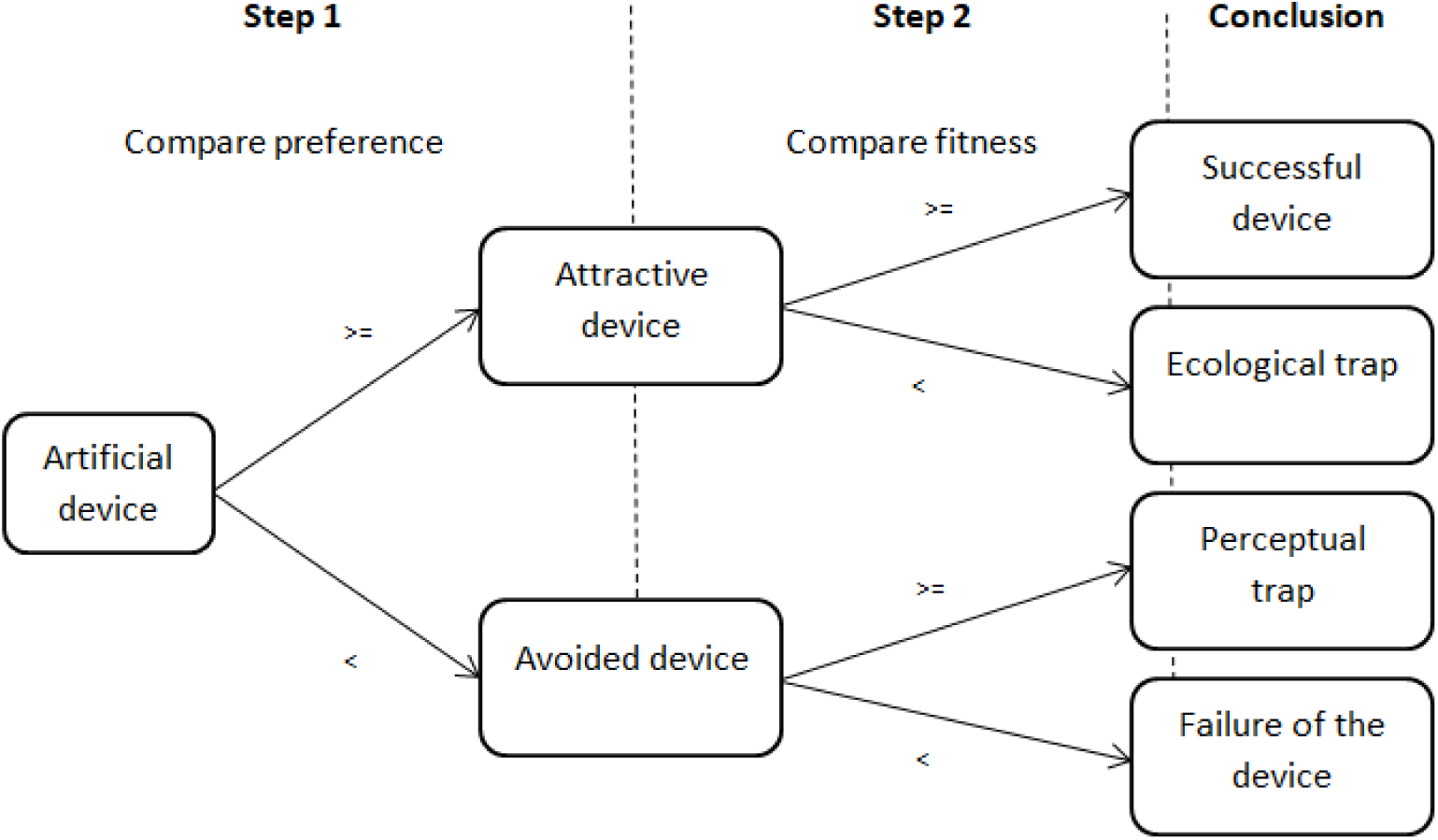
Conceptual framework for evaluating artificial conservation devices based on the ecological trap hypothesis: “>=” means values for artificial device are higher or equal compared to values for unmodified habitat; “<” means values for artificial device are lower compared to unmodified habitat.

Artificial shelters for cavity-using species, such as nest-boxes for birds or bats, are good models for testing the ecological trap hypothesis. They are probably the most widely used artificial devices for species conservation (Goldingay & Stevens, 2009). Their popularity among conservationists is mostly due to their low cost and the ease with which they can be deployed and surveyed (Hayward, Steinhorst & Hayward, 1992; Lambrechts et al., 2010). They are used in conservation programmes to respond to the growing disappearance of natural cavities in natural and semi-natural environments due to intensive forest management and landscape simplification (Lindenmayer et al., 2009). Numerous factors affect preference and fitness of species using cavities (see reviews in Goldingay & Stevens, 2009; Lambrechts et al., 2010), predation risk (Fontaine & Martin, 2006; Mönkkönen, Forsman, Kananoja, & Ylönen, 2009; Wesołowski, 2002; but see Lima, 2009; Martin, 1993; Pöysä, Ruusila, Milonoff, & Virtanen, 2001 for absence of effect on preference) and microclimate (Rhodes, O’Donnell, & Jamieson, 2009; Wachob, 1996) being among the most important for cavity selection. Temperature and humidity are key drivers of incubation and chick development, which can lead parents to select the most climatically favourable breeding environments (Rhodes, O’Donnell, & Jamieson, 2009). In artificial devices, the microclimate is not always as stable as in natural cavities; previous studies have shown that extreme temperatures in artificial boxes could impact the fitness of its occupants, e.g. in bats (Flaquer et al., 2006) or birds (Catry, Catry, Patto, Franco, & Moreira, 2015).

In this study, we took advantage of a dense population of European rollers *Coracias garrulus* (Fig. 1), breeding both in natural cavities and nest-boxes, to formally test the ecological trap hypothesis for an artificial conservation device. To do this, we compared occupation rates and breeding success of nest-boxes and natural cavities and investigated the impact of microclimate on breeding success and occupation probability.

The European roller (hereafter “the roller”) is an obligate secondary cavity breeder (Cramp, 1985): it does not create or modify its breeding site but uses already available sites on arrival after migration (Cramp, 1985). Thus, rollers respond very quickly to nest-box deployment (Aleman & Laurens, 2013) sometimes by abandoning nearby natural cavities in favour to nest-boxes (Valera, Václav, Calero-Torralbo, Martínez, & Veiga, 2019) and as such are very good models for studying nesting site preference. Due to this behaviour, we expected higher occupancy rates in nest-boxes compared to natural cavities. To our knowledge, no previous study compared occupancy between natural and artificial nests for rollers. We expected artificial nests to have less microclimatic buffering capacity than natural nests due to structural differences (Amat-Valero, Calero-Torralbo, Václav, & Valera, 2014; Catry et al., 2015; Maziarz, Broughton, & Wesołowski, 2017), and that these more variable conditions would lower breeding performance, following Catry, Catry, Patto, Franco, & Moreira, 2015 who found that roller chicks’ mass gain decreased with extreme heat events in nest-boxes but not in adobe wall cavities. Based on these two predictions, we expected nest-boxes to act as an ecological trap for the European roller.

## 2. Material and methods

### 2.1 The European roller

The European roller is a trans-Saharan migratory bird that winters in southern Africa (Finch et al., 2015) and breeds between May and July in a range that extends from the Mediterranean basin through Eastern Europe to central Asia (Cramp, 1985). Breeding sites are principally woodpecker cavities, but also include bee-eater burrows in sand cliffs and cavities in buildings such as farm houses or bridges (Cramp, 1985). Despite its recent re-ranking to “least concern” on the global IUCN Red List (IUCN, 2016), the European roller remains a threatened species in most European countries (Tokody et al., 2017). Rollers have benefited from numerous conservation programmes over the past two decades (Tokody et al., 2017), including the deployment of thousands of nest-boxes all over Europe (Finch et al., 2019).

### 2.2 Study site

The study took place in the valley of Les Baux-de-Provence (43°41’ N, 4°46’ E; WGS 84), located near the city of Arles, in the Bouches du Rhônes deparment of France. The region is classified as a meso-mediterranean bioclimatic stage, with hot and dry summers (average maximum temperature of 30°C in July). The valley extends over 2000 hectares of cereal fields and grasslands. A dense drainage channels’ network extending throughout the valley is bordered by several kilometres of hedges and riparian forest (Chambre d’Agriculture des Bouches du Rhône, 2008), where black poplars *Populus nigra* and white poplars *Populus alba* offer an exceptional density of natural cavities, created mostly by European green woodpeckers *Picus viridis* (Butler, 2001). This situation contrasts with other sites in southern France, where availability of nesting sites has been recently confirmed as a major limiting factor in roller breeding pairs’ density (Finch et al., 2019). Common starling *Sturnus vulgaris* is the main competitor of rollers for cavities in the study area, but breeds averagely earlier and is actively chased from the nesting sites by roller pairs at their arrival (A Rocha France, unpublished data).

### 2.3 Nest monitoring

We deployed 87 nest-boxes on private properties on the study site from 2002 to 2012 for the study and conservation of rollers, out of which 50 were still available between 2016 and 2019 (Fig. A1). The nest-boxes were built using plywood (15 mm thick) and had a volume of approximately 21,870cm^3^ (H1=25cm; H2=35cm; L=27cm; W=27cm) and 57-60 mm entrance hole. They were installed on trees at different heights and orientations. Nest-box occupation has been checked at least three times every year from mid-May to the end of June between 2016 and 2019, using a Voltcraft BS-250XIPSD endoscopic camera mounted on a telescopic pole. In breeding situations, at least three additional inspections were organized at c.a. one week intervals to survey the number of laid eggs, hatched eggs and fledglings per nest. All fledglings were ringed at between 15 and 25 days old. Fledging success was determined on the basis of the number of fledglings ringed, but was corrected retrospectively if a chick died after ringing and was found during the nest-box cleaning session. The date of the first egg-laying was back-calculated from nest monitoring observations (using two-days interval between the laying of each egg and 21 days between first egg laying date and hatching (Personnal field obs. and Guillaumot, 2016)). Nest abandonments and predation were also systematically recorded.

Between 2016 and 2019, a sample of accessible natural cavities located on the same study site was checked for occupation by rollers. Monitoring of the natural cavities was similar to that described for nest-boxes only in 2017. In other years, monitoring intensity was generally less frequent, and hence breeding parameters were only calculated for occupied nests visited at least three times between first egg-laying and first chick-fledging dates. For both nest types, roller breeding monitoring occurred between the 20 May and the 17 August.

In 2017, we explored the study site intensively and located 190 natural cavities suitable for roller reproduction on 154 different trees (Fig. A1). Suitability of cavities for roller reproduction was determined based on minimal hole diameter of occupied cavities at our study site (47mm) (A Rocha France, unpublished data) and location in the tree (no branches in front of the cavity hole enabling access for predators), all other tree and cavity charasterisitcs being not selected by rollers in the study site (see previous studies Butler, 2001; Dasse, 2016; Eltabet, 2013). For all occupied (n=23) and a sample of randomly selected unoccupied cavities (n=77) accessible by foot or with a 8m ladder, we measured height above ground, circumference of the tree at breast height (1.30m) and at cavity height, orientation, entrance height, entrance width, depth and length (Table 1).

**Table 1:** Name and description of the different microclimate and nest variables measured in natural cavities and nest-boxes in the Vallée des Baux (France). “%RH”: percentage of relative humidity.

|  | Name |  | Description |
| --- | --- | --- | --- |
|  | Daily variable | Mean variable over 15 consecutive days |  |
| Microclimate variables | MaxTint | MAXTint | Maximal temperature in the nest (°C) |
|  | MinTint | MINTint | Minimal temperature in the nest (°C) |
|  | MoyTint | MOYTint | Mean temperature in the nest (°C) |
|  | DiffTint | DIFFTint | Difference between maximal and minimal temperature in the nest (°C) |
|  | Dtmoyabs | DTMOYabs | Mean over a day of the absolute delta between interior and exterior temperature (°C) |
|  | Dtmaxabs | DTMAXabs | Absolute value of the daily maximum delta between interior and exterior temperature (°C) |
|  | Maxhum | MAXHum | Maximum humidity level in the nest (%RH) |
|  | Minhum | MINHum | Minimum humidity level in the nest (%RH) |
|  | Moyhum | MOYHum | Mean humidity level in the nest (%RH) |
|  | DiffHum | DIFFHum | Difference between maximal and minimal humidity level in the nest (%RH) |
| Nest variables | Name |  | Name of the nest-box or cavity |
|  | Type |  | Nest type : either nest-box or cavity |
|  | occupation |  | Occupation of the nest by rollers during the breeding season : 1 for roller nests, 0 otherwise |
|  | OccR |  | Real occupation of the nest during monitoring (roller or starling) : 1 for occupied nests sensor measurements, 0 for empty nests during sensor measurements |
|  | CH |  | Cavity height (m) |
|  | CBH |  | Circumference of the tree at breast height (1.30m) (cm) (natural cavities only) |
|  | CCH |  | Circumference of the tree at cavity height (cm) (natural cavities only) |
|  | Orientation |  | Entrance orientation of the nest (degrees) |
|  | Ori |  | Simplified orientation of the nest (either N, E, S or W with N: 316°-45°; E: 46°-135°; S: 136°-225°; W: 226°-315°) |
|  | EH |  | Entrance height (cavity) (mm) (natural cavities only) |
|  | EW |  | Entrance width (cavity) (mm) (natural cavities only) |
|  | CL |  | Cavity length (cm) (vertical) (natural cavities only) |
|  | CD |  | Cavity depth (cm) (horizontal) (natural cavities only) |
| Other variables | Date |  | Day of the measurement (Julian calendar date) |

### 2.4 Microclimate monitoring

In 2017, we monitored the microclimate in the nest-boxes (n=17) and natural cavities (n=23) occupied by rollers, as well as in a sample of randomly selected nest-boxes (n=16) and natural cavities (n=18) empty or occupied by Common starlings. Orientation and height of monitored nest-boxes did not differ from available nest-boxes (Orientation: Chi^2^=0.18, df=3, *P*=0.98; height: F-statistic=0.19, df=1,71, *P*=0.66). Monitored natural cavities had the same orientation but were in average 1.05m higher than the sample of measured available cavities (Orientation: Chi^2^=1.14, df=3, *P*=0.77; height: F-statistic=4.98, df=1,133, *P*=0.03) respectively. Interior temperature (°C) and humidity (percentage of relative humidity: %RH), and exterior temperature (°C) were respectively measured with iButtons DS1923 and DS1921G-F5 (Maxim Integrated Products, Inc.). Measurements were taken every hour for 15 consecutive days during the period between 6 June 2017 and 20 August 2017, which covers most of the incubation and chick rearing period in our study site (23 May to 10 August, based on 18 years of roller breeding monitoring on the study site, (A Rocha France, unpublished results). For occupied nests, sensors were deployed only after the clutch was complete, in order to prevent nest abandonment. In the nest-boxes, interior sensors were screwed directly to the rear wall, with the captor facing inside the box, and in the cavities, sensors were placed inside the hole at least 20 cm away from the entrance and held by a flexible aluminium angle bracket screwed directly into the bark of the tree. The aluminium rods were restricted to the sides of the cavity entrance in order to reduce possible disturbance of animal movements in and out of the cavity. Exterior sensors were placed in the shade, either under the base of the nest-boxes, facing the ground, or on the bark of the tree at cavity height, with the captor facing north. The occupation of each nest by rollers or Common starling (the only other species using the monitored nest-boxes and cavities during our study) during the measurement period was recorded.

The IButton data was extracted with OneWireViewer software (1-Wire drivers X86, version 4.03). The humidity data was corrected following the sensor manufacturer’s recommendations (Maxim Integrated Products Inc, 2011: p.53). For each nest and each day, we calculated: (1) the maximum, minimum and mean interior temperature and humidity rate, (2) the daily amplitude of the interior temperature and humidity (i.e. the difference between the minimum and maximum daily temperature/humidity rate inside the nest), (3) the daily mean and maximum absolute delta between the interior and exterior temperature (i.e. the mean and the maximum over 24 hours of the absolute values of hourly differences between exterior ambient temperature and interior nest temperature). Additionally, we calculated (4) the mean value over 15 consecutive days of each of the above parameters (Table 1).

## 3 Statistical analysis

### 3.1 Nest type preference

We compared the attractiveness of natural vs artificial nests in 2017 using the estimated occupancy rate (Johnson, 1980) and the first egg-laying date (Julian calendar date) as a proxy for settlement date of the rollers at nesting site (Robertson & Hutto, 2006). We considered only one cavity per tree in the sample (n=154), as rollers are territorial; to our knowledge more than one roller pair per tree has never been recorded. We compared occupancy rates between nest types using a Generalized Linear Model (GLM) with a binomial distribution and logit link function. However, as the sample included all nest-boxes (n=50) and occupied natural cavities (n=23) but only an unknown proportion of the available natural cavities (n=131) (inevitably lower than the actual amount), the true proportion of occupied natural cavities is lower than our estimate.

We tested the effect of nest type on the 2017 first egg-laying date using a Linear Model with nest type as the explanatory variable.

### 3.2 Breeding success

All the breeding success analysis were made for the 2016-2019 breeding seasons for all the occupied nests that were intensively monitored (n=69 and n=46 for nest-boxes and natural cavities respectively). We tested the difference of predation frequency (over breeding attempts with known outcome) between nest types using a Chi^2^ test. Successful nests were defined as roller nests with at least one fledgling. We tested the correlation between breeding success parameters and nest type using GLMs with nest type as the explanatory variable and four different breeding response variables (Poisson distribution and log link function for the number of eggs and number of fledglings per nest, binomial distribution and logit link function for the probability that a nest was successful and for the probability that an egg produce a fledging chick). Finally, for successful nests only (n=62 and n=25 for nest-boxes and natural cavities respectively), we tested the correlation of the probability that an egg produced a fledging chick, with nest type and egg-laying date, using a GLM with a binomial distribution and logit link function. We checked the overdispersion in the residuals when using a Poisson distribution and none was detected.

### 3.3 Microclimate parameters

All the nest characteristic variables were scaled prior to analysis. We tested the correlation of nest characteristic variables with microclimate parameters. We used Generalized Linear Mixed Models (GLMMs) with a Gaussian error structure, with the individual nest name (nest ID) and date (Julian calendar) as random factors. We compared cavity height and orientation between nest types, using a linear model with a Gaussian error structure and a Chi^2^ test, respectively. We tested the effect of nest height and orientation for both nest types on microclimate parameters. The models included the interaction of nest type (in order to control for nest characteristics disparities between nest types) and real occupation of the nest (i.e. nest occupation during sensor deployment) as additive factor (in order to control for the potentially induced modification of the microclimate (Maziarz, 2019; Vel’ký, Kaňuch, & Krištín, 2010)) (see ‘Results’). All the other nest characteristics were available only for natural cavities and hence their correlation with microclimate was tested solely on the subset of natural cavities, with real occupation of the nest as additive factor in the models.

We compared the daily microclimate variables between nest types, using GLMMs with a Gaussian error structure, with nest name and date as random factors. We included the interaction of real occupation in order to control for the response to the presence of birds in the nest according to each nest type.

### 3.4 Effect of microclimate on breeding success and occupation probability

We tested the correlations of the mean microclimate variables over 15 consecutive days with (1) the number of eggs, (2) the number of fledglings, (3) the probability that a nest was successful, (4) the probability that an egg produced a fledging chick and (5) the occupation probability by rollers, using GLMs with a Poisson and log link (1,2) and binomial distribution and logit link (3,4,5), respectively. For each microclimate variable, we controlled for possible disparities between nest types by including the interaction effect of nest type in the models. All statistical analysis were performed with R environment version 3.4.3 (R Core Team, 2017) using packages nlme (Pinheiro, Bates, DebRoy, Sarkar, & R Core Team, 2016), lme4 (Bates, Mächler, Bolker, & Walker, 2015) and lmerTest (Kuznetsova, Brockhoff, & Christensen, 2017).

## 4 Results

### 4.1 Nest type preference

Occupancy rate of nest-boxes was significantly higher than that of natural cavities (0.34 95% CI [0.20;0.52] and 0.15 95% CI [0.10;0.21] respectively, z= 2.87, df=1,203, *P*<0.01). The egg laying dates were similar for both nest types (146.6 95% CI [143.2;150.0] and 150.3 95% CI [145.3;155.2] for natural and artificial nests respectively) (F_1,35_=2.14, *P*=0.15).

### 4.2 Breeding success

The breeding parameter estimates are summarized in Table 2. No significant difference was found between nest types for the probability that a nest was successful (z=1.01, df=1,3,114, *P*=0.31) or for the probability that an egg produced one fledging chick in successful nests (z=−0.17, df=1,3,86, *P*=0.86). Nor was there a significant difference between nest types for average clutch size (z=0.21, df=1,3,101, *P*=0.84) or average number of fledglings (z= 1.30, df=1,3,106, *P*=0.19). 14 recorded breeding attempts failed during the 2016-2019 breeding seasons on the study site (7 in nest-boxes and 7 in natural cavities) out of which 10 were predated, with no significant difference in frequencies between nest types (4 in nest-boxes and 6 in natural cavities, Chi^2^=1.13, df=1, *P*=0.29).

**Table 2:** Estimates of breeding parameters for European rollers breeding in nest-boxes or cavities between 2016 and 2019 in the Vallée des Baux (France) with corresponding 95% confidence interval and sample size (n).

|  | Nest-boxes |  |  | Natural cavities |  |  | All nests |  |  |
| --- | --- | --- | --- | --- | --- | --- | --- | --- | --- |
|  | Estimate | 95% CI | n | Estimate | 95% CI | n | Estimate | 95% CI | n |
| Probability that a nest was successful | 0.76 | [0.50-0.91] | 69 | 0.64 | [0.37-0.84] | 46 | 0.72 | [0.52-0.86] | 115 |
| Probability that an egg produced a fledging chick | 0.75 | [0.62-0.85] | 62 | 0.74 | [0.56-0.87] | 25 | 0.75 | [0.63-0.84] | 87 |
| Average clutch size | 5.13 | [4.26-6.18] | 65 | 5.03 | [3.98-6.36] | 37 | 5.10 | [4.20-6.19] | 102 |
| Average number of fledglings | 3.88 | [3.09-4.88] | 69 | 3.71 | [2.66-5.17] | 38 | 3.86 | [2.95-5.04] | 107 |

### 4.3 Effect of nest characteristics and ambient temperature on microclimate

Monitored natural cavities were significantly higher than monitored nest-boxes (CH = 6.01m 95% CI [5.32;6.70] for natural cavities, CH=4.38m 95% CI [3.35;5.41] for nest-boxes) (F-statistic=9.65, df=1,61, *P*<0.01). The orientation of the nests monitored for microclimate differed between nest types (Chi^2^=11.34, df=3, *P*=0.01), with only one north-facing nest-box (3% *vs* 17% for natural cavities) and only two east-facing nest-boxes (7% *vs* 29% for natural cavities) (Fig. A2).

For both nest types, microclimate parameters were not correlated to nest height (Table A1.a). In natural cavities, western orientation had a slightly lower average and maximum temperature buffering capacity than southern orientation (Dtmoyabs: −1.43°C, *P*=0.06; Dtmaxabs: −2.90°C, *P*=0.05). Northern orientation had a slightly lower maximal temperature than southern and eastern (MaxTint: South: −1.99°C, *P*=0.07; East: −1.89°C, *P*=0.09) and a smaller interior temperature variation (DiffTint: South: −2.34°C, *P*=0.03; East: −2.27°C, *P*=0.04) (Fig. A3.A & B). Interior nest humidity was correlated to orientation, with north-facing nests more humid than those oriented towards other directions (Maxhum: East: +9.75%RH, Minhum: +17.95 to +21.12%RH, Moyhum: +12.11 to +14.41%RH), with less variation in humidity (DiffHum: −9.57 to −14.57%RH) (Table A1.b) (Fig. A3.C) (Table A1.b). In nest-boxes, we compared only the western and southern orientations (because of small sample size for northern (n=1) and eastern (n=2) orientations) and found no differences between the two, except for maximum and average buffering capacity, which was significantly lower in the southern direction (Dtmoyabs: −0.40°C; Dtmaxabs: −0.98°C) (Table A1.b).

For natural cavities only, maximum and average interior temperature decreased with the tree circumference at breast height (CBH) (MaxTint: β_x_=−0.85, SE=0.42, *P*=0.05; MoyTint: β_x_=−0.60, SE=0.33, *P*=0.08). Similarly, maximum interior temperature and interior temperature variation decreased with the tree circumference at cavity height (CCH) (MaxTint: β_x_=−0.80, SE=0.37, *P*=0.04; DiffTint: β_x_=−1.02, SE=0.35, *P*=0.01) (Fig. A4.A & B). Cavities in thicker trunks or branches tended to be more humid, with minimum and average humidity rates slightly increasing with CCH (Minhum: β_x_=4.92, SE=2.61, *P*=0.07; Moyhum: β_x_=3.71, SE=2.19, *P*=0.099) (Fig. A4.C) (Table A1.a).

### 4.4 Comparison of microclimate parameters between nest types and occupation status

Recorded interior and exterior temperatures for all nests ranged from 12.53°C to 43.45°C and 11.50°C to 45.50°C respectively (minimum and maximum values over the whole study period).

Empty nest-boxes had a higher maximum (MaxTint: +2.51°C, *P*<0.01), average (MoyTint: 1.14+°C, *P*=0.01) and lower minimum (MinTint: −1.50°C, *P*<0.01) interior temperature, higher temperature variations (DiffTint: +88%, *P*<0.01) and a lower average temperature buffering capacity (Dtmoyabs: −55%, *P*<0.01) than empty cavities. Empty nest-boxes also had a more variable (DiffHum: +47%, *P*<0.01) and lower humidity rate (Moyhum: −27.24%RH, *P*<0.01; Maxhum: −29.90%RH, *P*<0.01; Minhum: −27.87%RH, *P*<0.01) than empty cavities (Fig. 3A). These differences were still significant and similar when birds were present in the nest during the monitoring period, except for average interior temperature, which tended to be higher in occupied cavities than in occupied nest-boxes (MoyTint:+2.44°C, *P*<0.01) (Fig. 3B) (Table A2).

**Figure 3:**
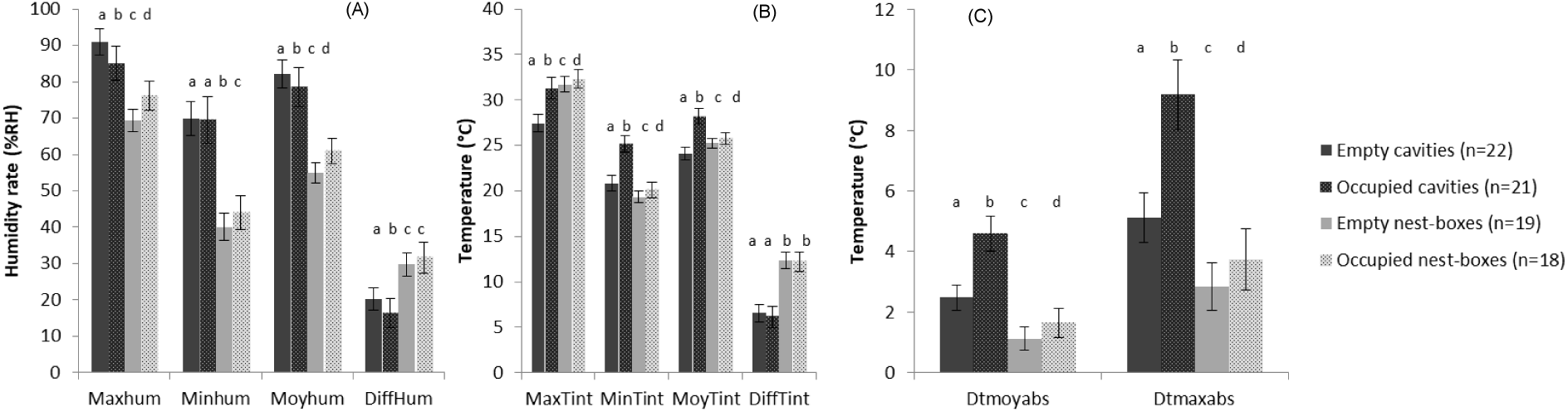
Comparison of the estimates of microclimate parameters (with their 95% confidence interval) between empty and occupied natural cavities and nest-boxes in the Vallée des Baux (France). (A) maximum, minimum, average and interior variation in humidity (Maxhum, Minhum, Moyhum and DiffHum), (B) maximum, minimum, average and interior variation in temperature (MaxTint, MinTint and MoyTint, DiffTint), and (C) maximum delta between interior and ambient temperature (Dtmaxabs) and average buffering capacity (Dtmoyabs). a, b, c & d: for each parameter, different letters indicate significant differences between the estimates of the corresponding nest type.

Maximum, average and minimum interior temperature were higher in occupied nests than in empty nests for both nest types (Nest-boxes: MaxTint:+0.6°C, *P*=0.06; MoyTint: +0.76°C, *P*<0.01; MinTint:+0.51°C,*P*=0.01 / Natural cavities: MaxTint:+3.82°C, *P*<0.01; MoyTint: +4.30°C, *P*<0.01; MinTint:+4.09°C,*P*<0.01). Similarly, occupation increased temperature buffering capacity in nests of both types (Nest-boxes: Dtmoyabs:+46%, *P*<0.01; Dtmaxabs: +32%, *P*=0.01; / Natural cavities: Dtmoyabs:+85%, *P*<0.01; Dtmaxabs: +80%, *P*<0.01) (Fig. 3C), but it did not modified interior temperature variation (Nest-boxes: DiffTint: −7%, *P*=0.68; / Natural cavities: DiffTint: −1 %, *P*=0.34). However, occupied nest-boxes tended to be more humid than empty ones (Moyhum: +6%RH, *P*<0.01; Maxhum: +6.70%RH, *P*<0.01; Minhum: +3.97%RH, *P*=0.01), while occupied cavities were significantly drier than empty cavities (Moyhum: −3.61%RH, *P*<0.01; Maxhum: −5.96%RH, *P*<0.01). Humidity rates were more stable in occupied natural cavities than in empty cavities (DiffHum: −19%, *P*=0.02) but they had similar daily fluctuations in empty and occupied nest-boxes (Table A2).

### 4.5 Effect of microclimate on occupation probability and breeding success

Roller occupation probability increased slightly with average interior temperature (MOYTint) in nest-boxes (*P*=0.08) but not in natural cavities (Fig. 4A). It increased marginally with maximum interior temperature for both nest types (MAXTint *P*=0.10) (Fig. 4B) but significantly with buffering capacity of the nest (DTMAXabs *P*=0.05 and DTMOYabs *P*<0.01) for nest-boxes only (Fig. 4C) (Table A3).

**Figure 4:**
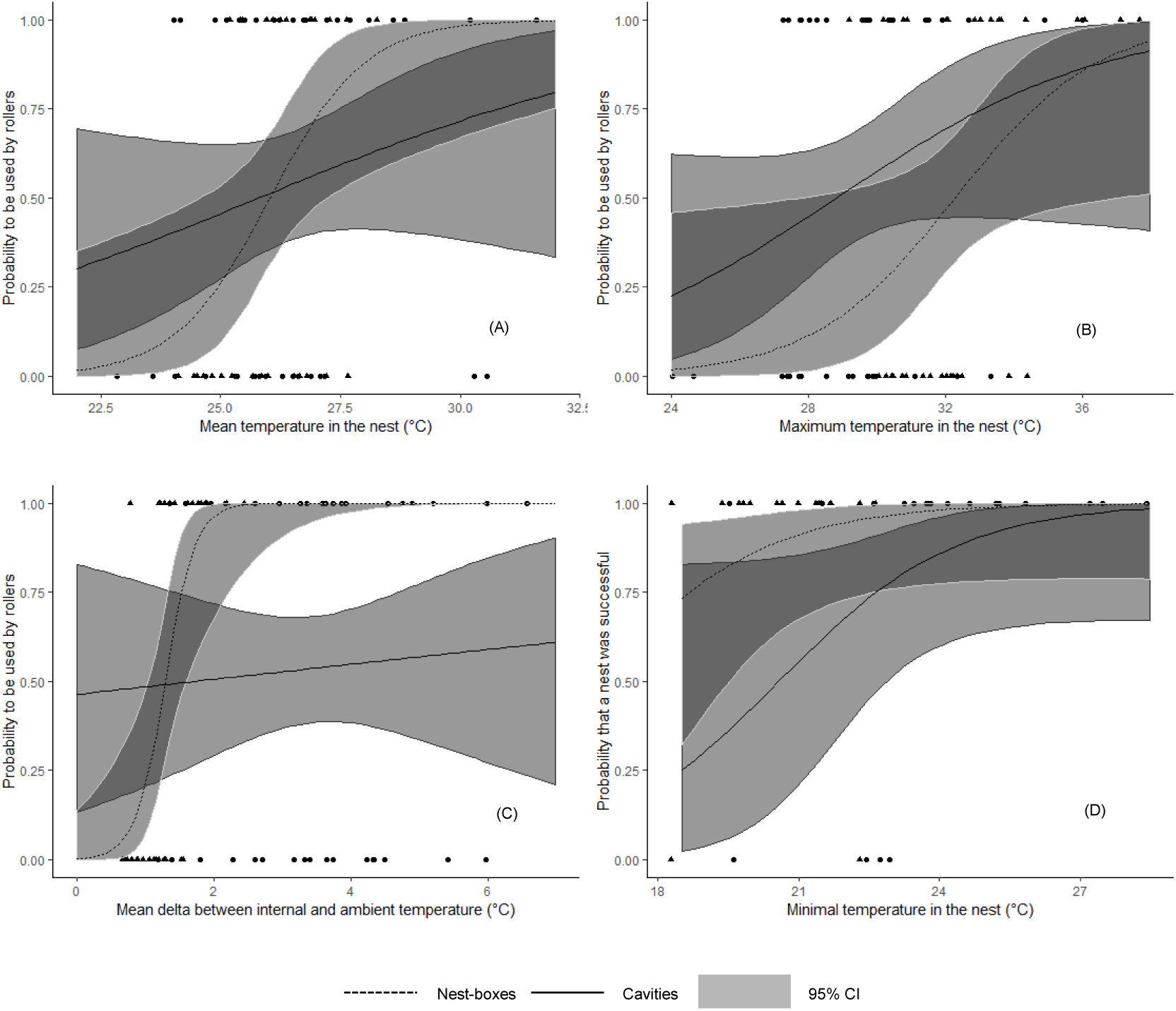
Correlation of different microclimate variables with occupation probability by the European roller (A: MOYTint, B: MAXTint and C: DTMOYABS) and with the probability that a roller nest was successful (D: MINTint) for nest-boxes (▴: dotted regression lines) and cavities (•: solid regression lines) in the Vallée des Baux (France). The 95% confidence intervals of regression lines are shown in grey, dark grey indicates overlaps between the intervals of the two nest types.

We found no effect of microclimate on clutch size, number of fledglings or fledging success per egg neither with nest type as an interaction factor (Table A3). However, in natural cavities, most microclimate variables were correlated with the probability that a nest was successful. This probability increased significantly with humidity rates (MOYhum *P*=0.02, MINhum *P*=0.02 and MAXhum *P*=0.03), slightly with average buffering capacity (DTMOYabs *P*=0.09) and minimum interior temperature (MINTint *P*=0.08) (Fig. 4D), whereas it decreased marginally when interior temperature and humidity variation increased (DIFFTint *P*=0.06 and DIFFhum *P*=0.07 respectively) (Table A3).

## 5 Discussion

Our results showed a strong contrast between nest types in terms of microclimatic conditions. Natural cavities had higher humidity on average and buffered ambient temperature much better than nest-boxes. Our results showed an impact of different microclimate parameters on occupation probability by rollers, which selected the warmest and better buffered artificial nests. The higher occupation rates of nest-boxes and similar egg-laying dates showed that artificial devices were preferred or equally selected by rollers compared to natural cavities, but we did not find any significant differences in roller breeding success between the two nest types. This suggests that nest-boxes are not ecological traps for European rollers at this study site.

### 5.1 Microclimate differs significantly between nest types

Nest orientation was correlated with several microclimate parameters (Table A1.b), confirming other studies that showed that this nest characteristic is an important microclimatic driver in both cavities and nest-boxes (Ardia, Pérez, & Clotfelter, 2006). North-facing nests are less exposed to sunlight – and hence to heat – compared to other orientations, and thus tend to have lower interior maximum temperatures (Wiebe, 2001) and higher humidity rates (Di Maggio, Campobello, Mascara, & Sarà, 2015).

Due to the deployment strategy of the nest-boxes, they differ significantly from natural cavities in terms of nest height and orientation, with cavities being averagely 1.63m higher and nest-boxes being oriented mostly to the West or to the South (Fig. A2). These differences could act as confounding factors for potential microclimate differences between nest types.

However, nest height was not correlated to any microclimate parameter (Table A1.a). Furthermore, even after controlling for nest orientation, we found strong microclimate differences between nest types, both for temperature and humidity rate (Fig. 3). In contrast to natural nests, the interior temperature of nest-boxes was closely linked to ambient temperature: interior temperatures were on average less than 1.2°C above or below the ambient temperature in nest-boxes, compared to 2.5°C in natural nests. Average interior temperature variations were also twice as high in artificial nests compared to natural cavities. Hence, artificial nests had more than 4°C higher maximal and 1.5°C lower minimal temperatures compared to natural cavities. Furthermore, nest-boxes were dryer, with 27% lower average humidity rate, varying of over 47% more on average. This suggests that humidity inside nest-boxes was closely linked to ambient humidity rates, while natural cavities were better buffered. Taken together, these results confirm the poor temperature and humidity buffering capacity of plywood nest-boxes compared to natural tree cavities, as tree trunks provide much better climatic insulation compared to the thin walls of nest-boxes (Rowland, Briscoe, & Handasyde, 2017). Similar results were found in another study in the European roller between wooden nest-boxes and natural or semi-natural cavities in sand-cliffs, bridges and buildings (Amat-Valero et al., 2014; Catry et al., 2015). Combined with our results, this demonstrates that wooden nest-boxes are not reproducing the microclimate of any of the most common natural or semi-natural nest types of the European roller throughout its breeding range (Cramp, 1985), and thus cannot be treated as an equivalent substitute of natural nesting sites.

The presence of birds in the nest (both European roller and Common starling) during sensor deployment affected microclimate significantly, increasing interior temperature and temperature buffering capacity in both natural cavities and nest-boxes. However, we found similar differences between nest-types in both empty and occupied nests (Table A2). These results are in line with previous studies that found that even a much smaller bird, great tits *Parus major*, increased interior temperature and temperature buffering capacity in nest-boxes (Maziarz & Wesołowski, 2013; Vel’ký et al., 2010) as well as marsh tits *Poecile palustris* in natural cavities (Maziarz, 2019). Similarly to Maziarz 2019, and Maziarz and Wesołowski (2013), we found that humidity rates were lower in occupied compared to empty cavities (Table A2). However, we found a reverse relationship in nest-boxes. This result could be explained by the huge difference of humidity rates between nest types (Fig. 3): the presence of birds in the nest could buffer the humidity level to a medium level, in between the dry empty nest-boxes and the humid empty cavities. The effect of the presence of birds in the nest on microclimate underlines the greater energetic cost for birds using nest-boxes which showed greater microclimatic variations compared to natural cavities in our study site (Maziarz, 2019).

### 5.2 Rollers prefer artificial over natural nests despite more stressful microclimatic conditions

We found that nest-boxes had higher occupation rates compared to natural cavities, suggesting their strong attractiveness. This observed preference could be due to a better protection against predators in nest-boxes than in natural cavities, a benefit which has been identified for other species e.g. in the Madeiran storm petrel *Oceanodroma castro* (Bolton, Medeiros, Hothersall, & Campos, 2004). In our study, we found a slightly higher frequency of predation (excluding unknown failure events) in natural nests compared to nest-boxes (n=6 and n=4 respectively, 8.7% and 5.7% of recorded breeding attempts in respective nest type in our study site between 2016 and 2019). Although this difference is not significant on our study site, the reduced intensity in the monitoring of cavities compared to nest-boxes (see Methods) could mean that we could have not detected some predation events at the early stage of the reproduction (e.g. during egg-laying or incubation period). The real frequency of predation events in natural cavities might hence be higher than the one we observed. Long term breeding monitoring and more intensive surveys of natural cavities would give more insights to validate this hypothesis in the future.

Observed predation rates were however extremely low in both nest types and support the findings of Parejo and Avilés (2011) showing that rollers are able to assess predation risk in nest site selection, similarly to several cavity nesting birds (see e.g. Fontaine & Martin, 2006; Mönkkönen, Forsman, Kananoja, & Ylönen, 2009; Wesołowski, 2002). It would thus be possible that preference for safe nesting sites could lead rollers to select for less suitable thermal or humidity conditions in order to reduce the nest predation risk, as suggested in the great tit (Maziarz, Wesołowski, Hebda, Cholewa, & Broughton, 2016). However, this hypothesis is not supported by our findings for the roller: Of the available nest-boxes, rollers selected the warmest ones, with high maximal (MAXTint) and average interior temperature (MOYTint) (Fig.4A & 4B) (Table A3). Warmer nests may be especially attractive to cavity-nesting altricial birds, as they could reduce energetic costs and improve hatching success, especially at the beginning of the breeding season, when minimal ambient temperatures are still low (Reid, Monaghan, & Ruxton, 2000). Early-season microclimatic conditions in nest-boxes could lead rollers to select these warmer artificial nests at the risk of exposing their chicks to more stressful extremely high temperatures in the following weeks (Catry et al., 2015). However, when selecting a nest-box, rollers also tended to occupy the nests with better temperature buffering capacity (DTMAXabs and DTMOYabs) (Fig. 4C) (Table A3). This result suggests that rollers could be able to avoid the most stressful microclimatic conditions among the available nest-boxes, independently from nest predation risk.

### 5.3 Roller breeding success does not differ between nest types

Contrary to our prediction, we found no significant difference in breeding success parameters between artificial and natural nests (Table 2).

The probability of a breeding pair bringing at least one chick to fledging in occupied natural nests was significantly positively correlated with humidity rates (MAXhum, MINhum, MOYhum), marginally negatively correlated with humidity and temperature variation (DIFFHum, DIFFTint) and marginally positively correlated with buffering capacity (DTMOYabs) and minimum temperature (MINTint) in the nest (Table A3). These results confirm that dry and poorly buffered microclimatic conditions negatively impact fitness.

However, all these parameters were significantly more favourable in natural cavities than in nest-boxes (Table A2). Thus we would have expected higher breeding success in natural nests, as demonstrated in other bird species, e.g. in the African penguin *Speniscus demersus* (Lei, Green, & Pichegru, 2014) or in the Lesser kestrel *Falco naumanii* (Catry et al., 2015). The absence of significant disparities between nest types could be explained by roller occupation strategy. As rollers avoided the nest-boxes with the least favourable microclimates (see section 5.2), the induced stress in the selected artificial nests may have been sufficiently reduced as to not significantly impact the individual fitness. The positive impact of this strategy may have been additionally strengthened by the particular heat tolerance of rollers compared to other bird species. Catry et al. (2015) found that in nest-boxes in Portugal roller chicks suffer much less from extreme heat conditions than those of the Lesser kestrel. It is possible that rollers have sufficient plasticity to tolerate a certain amount of microclimatic stress. Furthermore, Catry et al. (2015) showed that roller chicks could survive interior maximum temperatures of more than 50°C, which largely exceeds maximum interior temperatures recorded in our study (43.5°C), despite similar heat waves with days exceeding 37°C of maximum exterior temperature in June, July and August 2017 (www.meteofrance.fr). Nest-boxes directly exposed to sunlight can have interior maximum temperature that largely exceeds ambient temperature (Ellis, 2016). All the nest-boxes in our study area are placed on tree trunks and therefore benefit from the shade of branches and foliage, which is not the case in nest-boxes placed on poles and buildings in the study site described by Catry et al. (2015). This could explain the lower maximum temperatures found in the nest-boxes on our study site.

In southern Spain, Valera et al. (2019) also did not find breeding success disparities in the European roller between wooden nest-boxes and natural and semi-natural cavities, despite microclimatic disparities between nest types similar than found in our study (Amat-Valero et al., 2014). However, to our knowledge, they did not explore the link between the two. Overall, our results found that roller fitness was not negatively impacted in the nest-boxes they selected, at least in terms of breeding success. However, the least favourable microclimate in nestboxes probably increases energy expenditure of rollers, as also suggested in the great tit (Maziarz, 2019), which could have other fitness consequences, and affect traits such as chick growth (Andreasson, Nord & Nilsson, 2018), body condition (Catry, Franco et al., 2011), or post-fledging survival (Greño, Belda & Barba, 2008), which would be valuable to explore in future studies. Nonetheless, according to our conceptual framework (Fig. 2), in our study site, nest-boxes for rollers must be considered successful devices for reproduction. By avoiding the nest-boxes with the worst microclimate, it is possible that rollers avoid potential ecological traps, contrary to our initial hypothesis. It would be necessary to test this hypothesis in the future by comparing the influence of microclimate on occupancy in the roller between artificial and natural nests, over a climatic and geographical gradient.

### 5.4 Recommendations for nest-boxes placement

In southern France, roller breeding density is limited by nest availability (Finch et al., 2019). In our study site, nest-boxes were more attractive than natural cavities and did not deter breeding success of rollers. Therefore, providing nest-boxes as additional available nesting places could be beneficial to conservation of the species.

While we found that rollers seem to be able to avoid the least favourable nest-boxes among those available, this choice was only possible because these were located in different settings, under tree canopy, which created a variety of microclimates. This might not always be the case: for instance, in open habitats, nest-boxes are usually placed on electricity poles <u>(see e.g.</u> Aleman & Laurens, 2013; Monti et al., 2019) and hence directly exposed to sunlight. To avoid extreme microclimatic conditions in nest-boxes for cavity breeders such as rollers, we recommend the following:

1. In Mediterranean and tropical regions, place nest-boxes in shaded conditions and oriented northwest to northeast (southwest to southeast in Southern hemisphere). This reduces both maximum temperatures and temperature variation (see also Catry, Franco & Sutherland, 2011). However, in colder regions of the northern hemisphere, nest-boxes should be south-facing in order to favour early-breeding species, especially at the beginning of the breeding season (see e.g. Ardia et al., 2006; Dawson, Lawrie, & O’Brien, 2005). Similarly, we recommend avoiding placing nest-boxes on electricity or telephone poles, as these provide no shade. This is especially important in warm regions, where birds can suffer from overheating, but cold nights can also affect bird survival in every part of the world in such poorly buffered microclimatic conditions.
2. Build nest-boxes using materials with good insulation properties, such as carved logs or tree trunks (Griffiths et al., 2018), as it will reduce maximal and increase minimal temperature in the nest, as well as daily fluctuations, and increase humidity rate, and hence improve attractivity of the nest and the probability of laid eggs to produce fledglings.

These measures should reduce the probability of nest-boxes creating deleterious microclimatic conditions for their occupants. Our findings also confirm previous studies showing that nest boxes poorly replicate the microclimate of tree cavities. Therefore, we highlight the need to focus on protecting cavity trees, while treating nest boxes as temporary solution in species conservation, whose significant installation and maintenance costs could be difficult to sustain in the long term (Goldingay, Thomas, & Shanty, 2018; Lindenmayer et al., 2017, 2009).

## 6 Conclusion

Our findings demonstrated that for the European roller in our study site, nest-boxes were more attractive than natural cavities and that breeding success between the two nest types was not significantly different. The rollers seemed to be able to avoid the least favourable nest-boxes, and while the microclimatic conditions of artificial nests were more variable, this neither deterred occupation nor translated into lower breeding success. However, the large microclimate differences observed between nest types might affect energy expenditure of birds and could hence deter other bird fitness parameters or proxies which would be valuable to explore in future studies.

Using the framework of the ecological trap hypothesis enabled us to compare preference and fitness between a natural habitat – tree cavities – and nest-boxes in order to evaluate the effectiveness of the latter for the conservation of a secondary cavity breeder. This framework is potentially applicable to any species in which some populations use restored or artificial habitats as well as unmodified or natural habitats at a given stage of their life history.

However, this framework focuses solely on the ecology of the target species, while evaluating habitat restoration programs should take other considerations into account. A more holistic approach could include an evaluation of (i) the durability or resilience of the restored or artificial habitat in comparison to the natural or unmodified habitat, (ii) the social and economic costs of restoration compared to conservation outcomes, and (iii) ecosystem functionality, i.e. placing restored habitats and target species in a broader ecological network, taking into account the relationship between landscape elements and their structuration, and the fitness of the target species, which would enable finer-scale recommendations.

## 7 Data Accessibility

*All data will be archived in appropriate long term and public data repository (e.g. Dryad)*

## 8 Author’s contribution

TS and AB conceived the ideas and designed methodology; TS and AG collected and analysed the data; TS and AB led the writing of the manuscript. All authors contributed critically to the drafts and gave final approval for publication.

## 9 Acknowledgements

We are grateful to the numerous volunteers and interns from A Rocha France who participated in data collection: Anthony Bahuaud, Mélanie Dasse, Lisa Gili, Armelle Hege, Alice Goerger, Vincent Robert, Romain Maillet and Pierre Paulhiac. We thank the private landowners and farmers, who have given access to their land for this study.

## Appendix A Supplementary figures and tables

**Figure A1:**
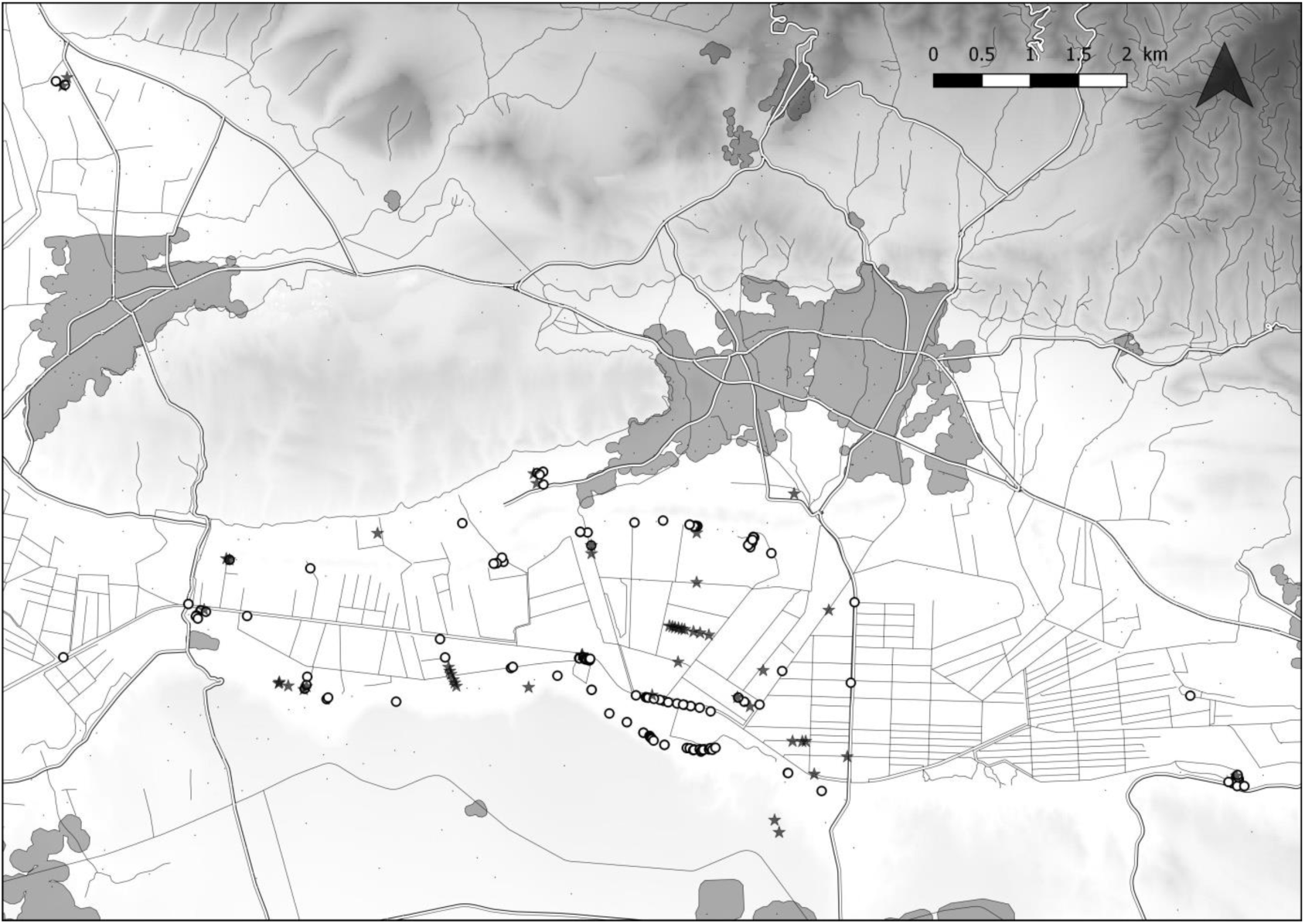
Map of the study area and distribution of natural cavities (rounds) and nest-boxes (stars) available for the reproduction of European rollers between 2016 and 2019.

**Figure A2:**
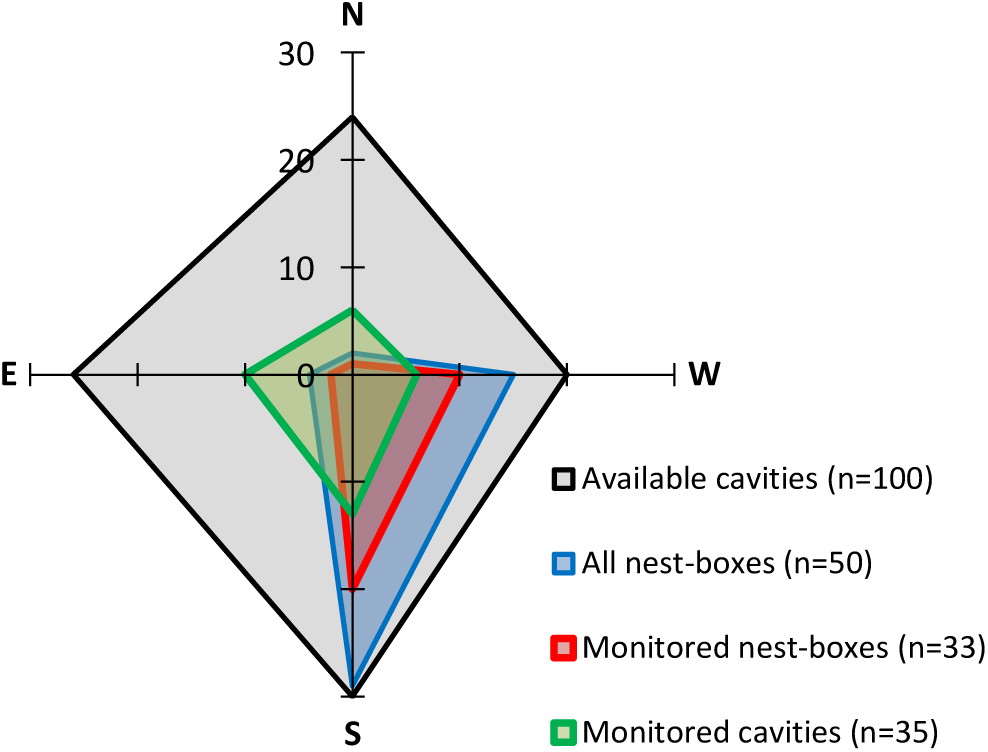
Orientation of cavities (n=35) and nest-boxes (n=33) monitored for microclimate, and of available cavities (n=100) and nest-boxes (n=50) for European roller reproduction in the Vallée des Baux (France) in 2017 (N: north, W: west, S: south, E: east).

**Figure A3:**
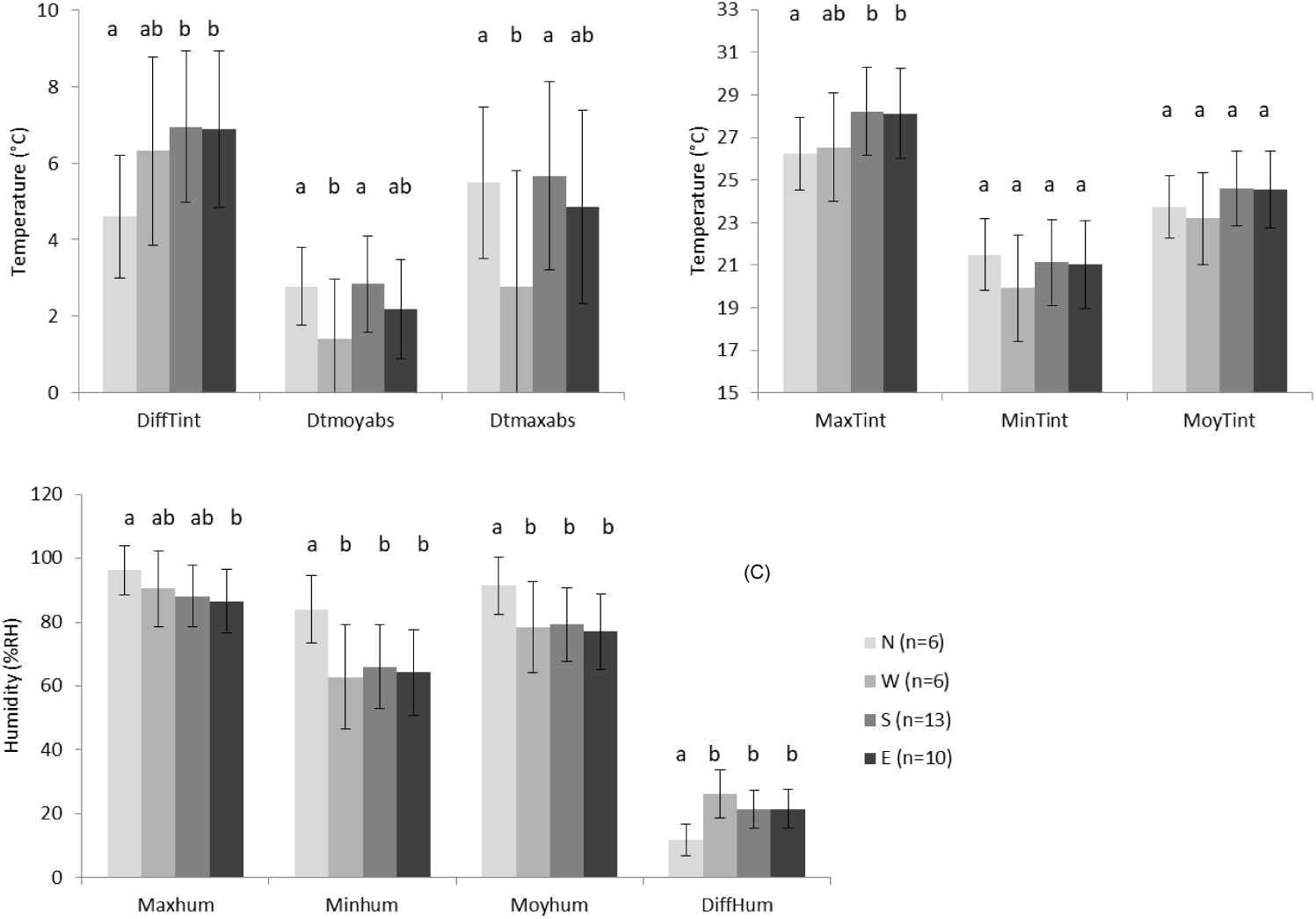
Estimates of microclimate parameters with their 95% confidence interval for different orientations (N: north, W: west, S: south, E: east) in natural cavities in the Vallée des Baux (France): (A) internal temperature variation (DiffTint), average buffering capacity (Dtmoyabs) and maximum delta between interior and ambient temperature (Dtmaxabs), (B) maximum, minimum and average interior temperature (MaxTint, MinTint and MoyTint respectively) and (C) maximum, minimum, average and interior variation in humidity (Maxhum, Minhum, Moyhum and DiffHum). a, b, c & d: for each parameter, different letters indicate significant differences between the estimates of the corresponding nest type.

**Figure A4:**
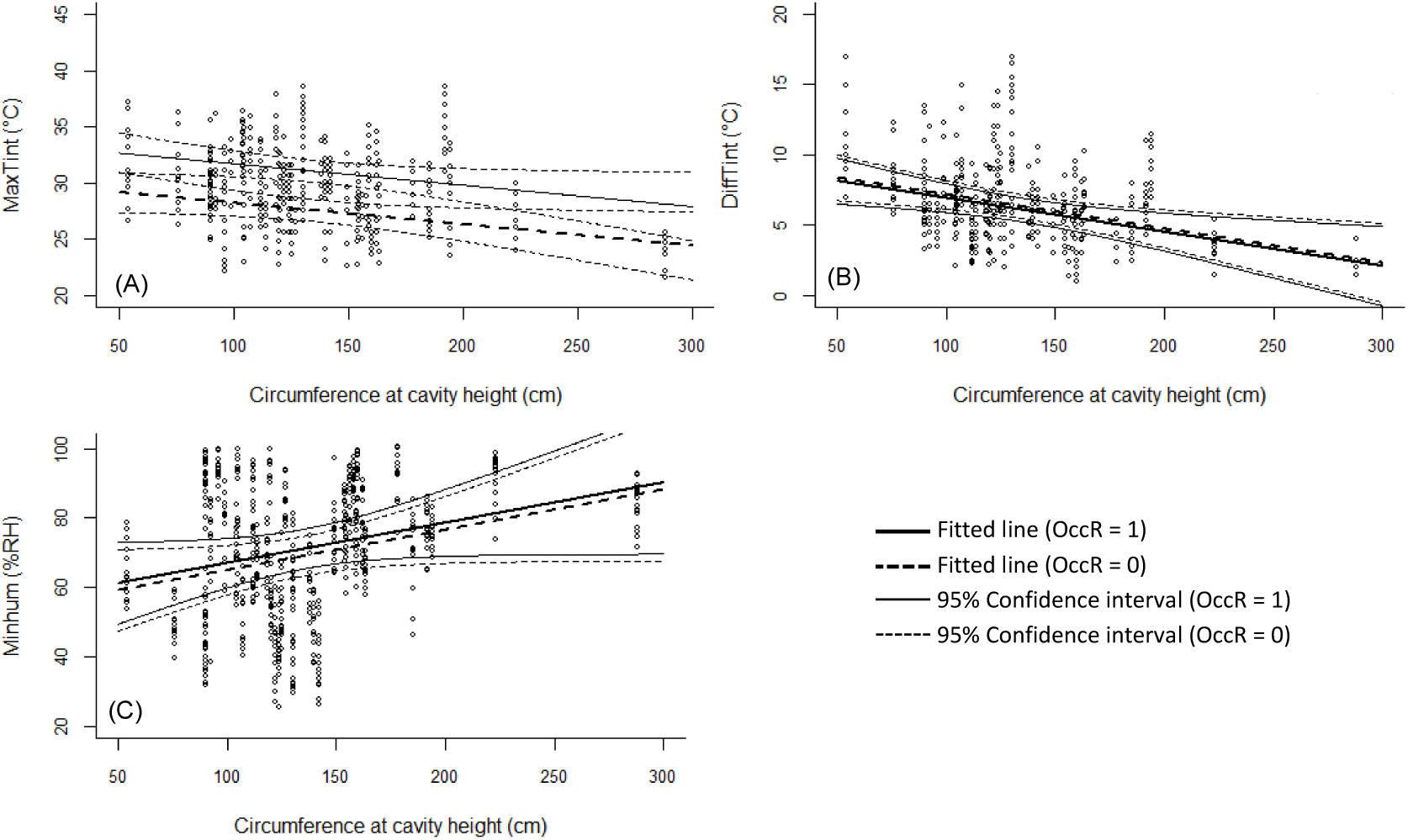
Correlation between circumference at cavity height and microclimate parameters in cavities available for European roller reproduction in the Vallée des Baux (France) with the additive effect of the occupation of nest during microclimate measurements (OccR): (A) maximum internal temperature (MaxTint); (B) internal temperature variation (DiffTint); (C) minimum humidity rate (Minhum). Dotted lines represent the 95% CI of the regression line.

**Table A1.a:**
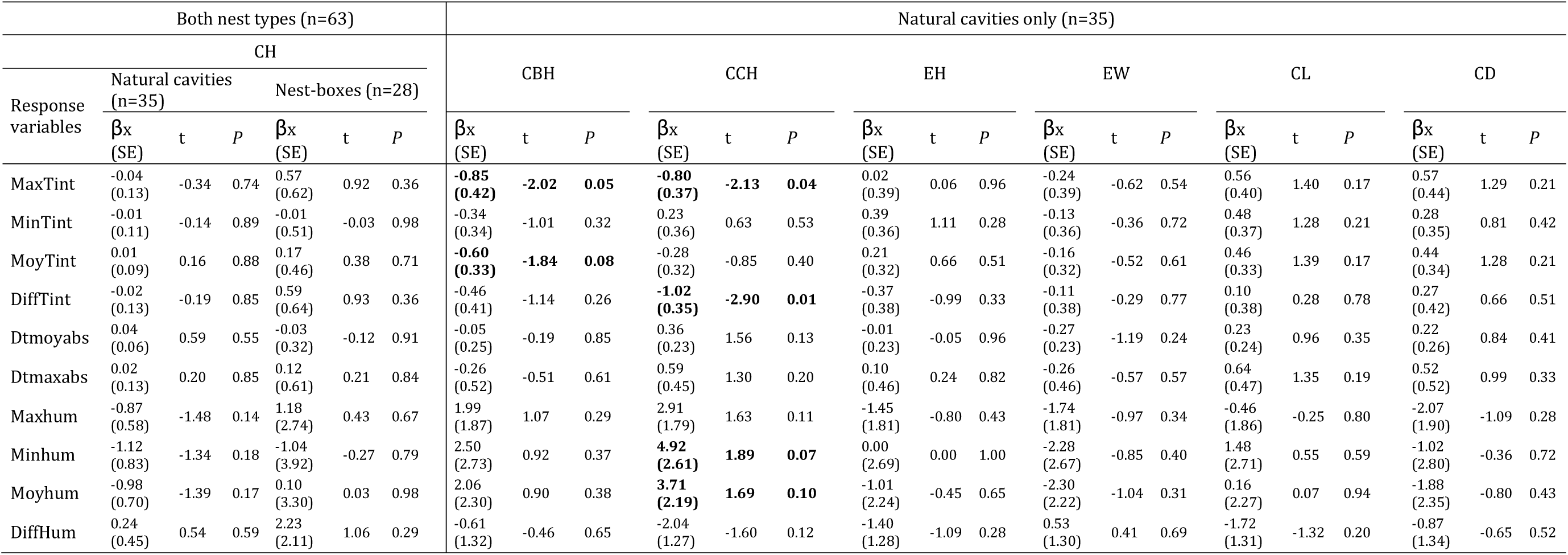
Correlations between nest characteristics and microclimate variables in nest-boxes and natural cavities available for European roller reproduction in the Vallée des Baux (France). CH: cavity height. CBH: circumference at breast height. CCH: circumference at cavity height. EH: entrance height. EW: entrance width. CL: cavity length. CD: cavity depth (=horizontal length). MaxTint: daily maximal temperature in the nest. MinTint: daily minimal temperature in the nest. MoyTint: daily mean temperature in the nest. DiffTint: difference between daily maximal and minimal temperature in the nest. Dtmaxabs: absolute value of the daily maximum delta between interior and exterior temperature. Dtmoyabs: mean over a day of the absolute delta between interior and exterior temperature. Maxhum: daily maximum humidity level in the nest. Minhum: daily minimum humidity level in the nest. Moyhum: daily mean humidity level in the nest. DiffHum: difference between daily maximal and minimal humidity level in the nest. “β_x_”: estimate of the slope of the correlation. “SE”: standard error of the estimate. “t”: t-value for the corresponding estimate. “*P”*: p-value. “n”: sample size. *P*-values lower than 0.10 are in bold.

**Table A1.b:**
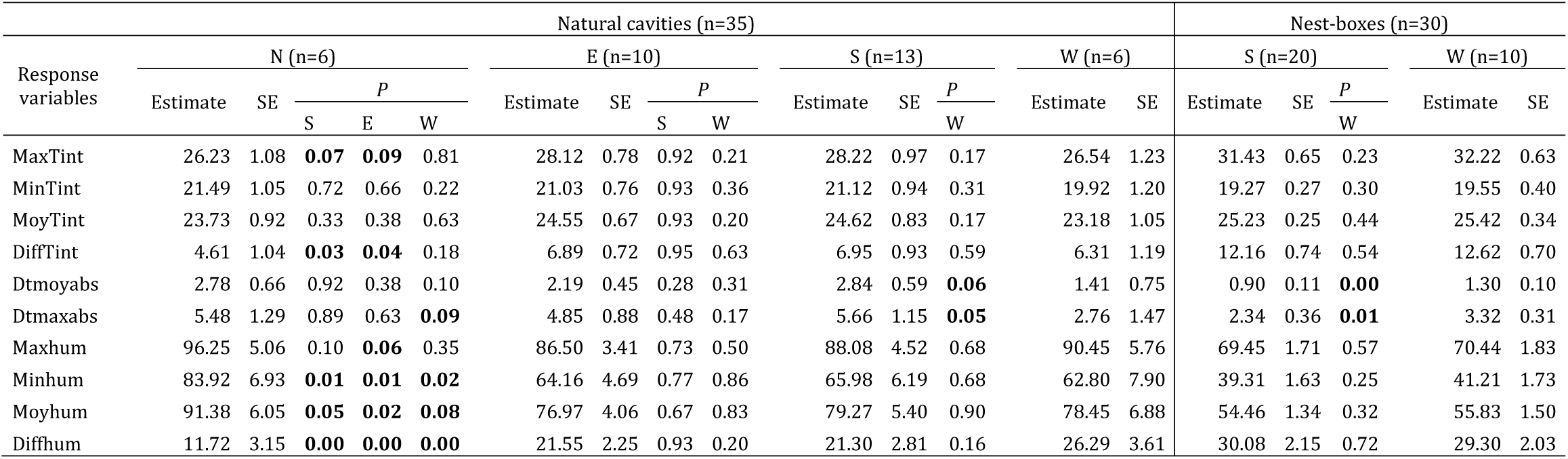
Comparison of the estimates of microclimate variables between the main orientations (N: north, W: west, S: south, E: east) of natural and artificial nests available for European roller reproduction in the Vallée des Baux (France). MaxTint: daily maximal temperature in the nest. MinTint: daily minimal temperature in the nest. MoyTint: daily mean temperature in the nest. DiffTint: difference between daily maximal and minimal temperature in the nest. Dtmaxabs: absolute value of the daily maximum delta between interior and exterior temperature. Dtmoyabs: mean over a day of the absolute delta between interior and exterior temperature. Maxhum: daily maximum humidity level in the nest. Minhum: daily minimum humidity level in the nest. Moyhum: daily mean humidity level in the nest. DiffHum: difference between daily maximal and minimal humidity level in the nest. “SE”: standard error of the estimate. “*P”*: p-value. “n”: sample size. *P*-values lower than 0.10 are in bold.

**Table A2:** Effect of cavity type and nest occupation during monitoring on different daily microclimate variables, in natural cavities and artificial nest-boxes available for European roller reproduction in the Vallée des Baux (France). MaxTint: daily maximal temperature in the nest. MinTint: daily minimal temperature in the nest. MoyTint: daily mean temperature in the nest. DiffTint: difference between daily maximal and minimal temperature in the nest. Dtmaxabs: absolute value of the daily maximum delta between interior and exterior temperature. Dtmoyabs: mean over a day of the absolute delta between interior and exterior temperature. Maxhum: daily maximum humidity level in the nest. Minhum: daily minimum humidity level in the nest. Moyhum: daily mean humidity level in the nest. DiffHum: difference between daily maximal and minimal humidity level in the nest. “SE”: standard error of the estimate. “t”: t-value for the corresponding estimate. “*P”*: p-value. “n”: sample size. *P*-values lower than 0.10 are in bold.

| Response variable | Empty cavities (n=22) |  | Empty nest-boxes (n=19) |  | Occupied cavities (n=21) |  | Occupied nest-boxes (n=18) |  | Paired t-tests |  |  |  |  |  |  |  |
| --- | --- | --- | --- | --- | --- | --- | --- | --- | --- | --- | --- | --- | --- | --- | --- | --- |
|  |  |  |  |  |  |  |  |  | Empty nest-boxes and cavities |  | Occupied nest-boxes and cavities |  | Occupied and empty nest-boxes |  | Occupied and empty cavities |  |
|  | Estimate | SE | Estimate | SE | Estimate | SE | Estimate | SE | t | P | t | P | t | P | t | P |
| MaxTint | 27.47 | 0.49 | 31.74 | 0.59 | 31.29 | 0.41 | 32.34 | 0.51 | 7.23 | **<0.01** | 1.77 | **0.08** | 1.87 | **0.06** | 9.28 | **<0.01** |
| MinTint | 20.85 | 0.43 | 19.35 | 0.48 | 25.15 | 0.34 | 20.11 | 0.42 | -3.13 | **<0.01** | -10.48 | **<0.01** | 2.89 | **<0.01** | 12.71 | **<0.01** |
| MoyTint | 24.11 | 0.38 | 25.25 | 0.42 | 28.20 | 0.26 | 25.76 | 0.32 | 2.71 | **0.01** | -5.78 | **<0.01** | 2.57 | **0.01** | 15.74 | **<0.01** |
| DiffTint | 6.56 | 0.47 | 12.36 | 0.60 | 6.13 | 0.44 | 12.21 | 0.56 | 9.63 | **<0.01** | 10.02 | **<0.01** | -0.41 | 0.68 | -0.96 | 0.34 |
| Dtmoyabs | 2.48 | 0.21 | 1.12 | 0.30 | 4.60 | 0.20 | 1.64 | 0.25 | -4.60 | **<0.01** | -9.92 | **<0.01** | 3.29 | **<0.01** | 10.73 | **<0.01** |
| Dtmaxabs | 5.11 | 0.42 | 2.85 | 0.59 | 9.18 | 0.41 | 3.75 | 0.52 | -3.86 | **<0.01** | -9.15 | **<0.01** | 2.74 | **0.01** | 10.00 | **<0.01** |
| Maxhum | 91.00 | 1.80 | 69.42 | 2.40 | 85.04 | 1.59 | 76.12 | 1.99 | -8.99 | **<0.01** | -3.69 | **<0.01** | 5.38 | **<0.01** | -3.74 | **<0.01** |
| Minhum | 69.96 | 2.40 | 40.03 | 3.30 | 69.45 | 1.90 | 44.00 | 2.37 | -9.07 | **<0.01** | -7.66 | **<0.01** | 2.72 | **0.01** | -0.27 | 0.79 |
| Moyhum | 82.21 | 1.99 | 54.97 | 2.73 | 78.60 | 1.47 | 60.97 | 1.82 | -9.99 | **<0.01** | -6.43 | **<0.01** | 5.38 | **<0.01** | -2.46 | **0.01** |
| DiffHum | 20.21 | 1.56 | 29.71 | 2.03 | 16.32 | 1.67 | 31.59 | 2.14 | 4.68 | **<0.01** | 7.42 | **<0.01** | 1.33 | 0.18 | -2.32 | **0.02** |

**Table A3:** Estimates of the interaction effect of mean microclimate variables over 15 consecutive days with nest type (natural cavities or artificial nest-boxes) on: breeding parameters (number of fledglings, clutch size, probability that a nest was successful and probability that an egg produced a fledging) and probability of a nest being used by rollers in the Vallée des Baux (France). MAXTint: mean of the daily maximal temperature in the nest. MINTint: mean of the daily minimal temperature in the nest. MOYTint: mean of the daily mean temperature in the nest. DIFFTint: mean of the difference between daily maximal and minimal temperature in the nest. DTMAXabs: mean of the absolute value of the daily maximum delta between interior and exterior temperature. DTMOYabs: mean of the mean over a day of the absolute delta between interior and exterior temperature. MAXHum: mean of the daily maximum humidity level in the nest. MINHum: mean of the daily minimum humidity level in the nest. MOYHum: mean of the daily mean humidity level in the nest. DIFFHum: mean of the difference between daily maximal and minimal humidity level in the nest. “n”: number of nest-boxes. “N”: number of natural cavities. “SE”: standard error of the estimate. “Z”: z-value for the corresponding estimate. “*P”*: p-value. “n”: sample size. *P*-values lower than 0.10 are in bold. a1, a2, b1 and b2 are the coefficients of the model equation y = a_1_ + a_2_ * W + b_1_ * X + b_2_ * X * W, for any response variable “y” and any microclimate variable “X” with W = 0 for natural cavities and W =1 for artificial nest-boxes.

|  |  | Response variables |  |  |  |  |  |  |  |  |  |  |  |  |  |  |  |  |  |  |  |
| --- | --- | --- | --- | --- | --- | --- | --- | --- | --- | --- | --- | --- | --- | --- | --- | --- | --- | --- | --- | --- | --- |
| Explanatory variables | Model coefficients | Number of fledglings (n=15; N=16) |  |  |  | Clutch size (n=15; N=16) |  |  |  | Probability that a nest was successful (n=17; N=20) |  |  |  | Probability that an egg produced a fledgling (n=15; N=16) |  |  |  | Probability of use by rollers (n=33; N=41) |  |  |  |
|  |  | Estimate | SE | Z | p | Estimate | SE | Z | p | Estimate | SE | Z | p | Estimate | SE | Z | p | Estimate | SE | Z | p |
| MAXTint | a1 | 1.51 | 0.15 | 10.39 | <0.01 | 1.64 | 0.14 | 11.99 | <0.01 | 1.17 | 0.59 | 1.98 | 0.05 | 2.02 | 0.46 | 4.40 | <0.01 | -7.38 | 4.56 | -1.62 | 0.11 |
|  | b1 | 0.17 | 0.18 | 0.96 | 0.34 | 0.11 | 0.17 | 0.66 | 0.51 | -0.64 | 0.65 | -0.98 | 0.33 | 0.51 | 0.56 | 0.92 | 0.36 | 0.26 | 0.16 | 1.65 | **0.10** |
|  | a2 | -0.13 | 0.21 | -0.63 | 0.53 | -0.05 | 0.20 | -0.27 | 0.78 | 1.66 | 1.45 | 1.14 | 0.26 | -0.58 | 0.58 | -1.00 | 0.32 | -8.10 | 8.73 | -0.93 | 0.35 |
|  | b2 | -0.24 | 0.24 | -1.00 | 0.32 | -0.17 | 0.22 | -0.77 | 0.44 | -0.29 | 1.19 | -0.25 | 0.81 | -0.57 | 0.66 | -0.85 | 0.39 | 0.22 | 0.28 | 0.80 | 0.42 |
| MINTint | a1 | 1.39 | 0.18 | 7.76 | <0.01 | 1.59 | 0.16 | 9.85 | <0.01 | 0.94 | 0.64 | 1.47 | 0.14 | 1.54 | 0.41 | 3.76 | <0.01 | 0.05 | 0.38 | 0.13 | 0.90 |
|  | b1 | 0.05 | 0.14 | 0.36 | 0.72 | 0.00 | 0.13 | 0.00 | 0.99 | 1.49 | 0.86 | 1.73 | **0.08** | 0.32 | 0.35 | 0.92 | 0.36 | 0.17 | 0.36 | 0.47 | 0.64 |
|  | a2 | -0.06 | 0.29 | -0.22 | 0.83 | -0.06 | 0.26 | -0.22 | 0.83 | 1.61 | 1.53 | 1.06 | 0.29 | -0.07 | 0.66 | -0.10 | 0.92 | 0.72 | 0.70 | 1.03 | 0.30 |
|  | b2 | -0.06 | 0.31 | -0.18 | 0.86 | -0.03 | 0.28 | -0.10 | 0.92 | -0.78 | 1.63 | -0.48 | 0.63 | -0.22 | 0.72 | -0.30 | 0.76 | 0.81 | 0.74 | 1.09 | 0.27 |
| MOYTint | a1 | 1.41 | 0.14 | 10.08 | <0.01 | 1.58 | 0.13 | 12.43 | <0.01 | 1.38 | 0.57 | 2.42 | 0.02 | 1.70 | 0.35 | 4.90 | <0.01 | -5.68 | 4.72 | -1.21 | 0.23 |
|  | b1 | 0.07 | 0.10 | 0.69 | 0.49 | 0.03 | 0.09 | 0.33 | 0.74 | 0.35 | 0.53 | 0.66 | 0.51 | 0.28 | 0.28 | 0.98 | 0.33 | 0.22 | 0.18 | 1.24 | 0.22 |
|  | a2 | -0.08 | 0.20 | -0.39 | 0.70 | -0.03 | 0.18 | -0.17 | 0.86 | 0.62 | 0.95 | 0.65 | 0.52 | -0.30 | 0.46 | -0.64 | 0.52 | -20.55 | 11.91 | -1.73 | **0.08** |
|  | b2 | -0.10 | 0.26 | -0.37 | 0.71 | -0.05 | 0.23 | -0.22 | 0.83 | -1.00 | 1.53 | -0.66 | 0.51 | -0.31 | 0.58 | -0.53 | 0.59 | 0.79 | 0.46 | 1.73 | **0.08** |
| DIFFTint | a1 | 1.57 | 0.26 | 60.3 | <0.01 | 1.75 | 0.24 | 7.27 | <0.01 | 0.02 | 0.90 | 0.02 | 0.98 | 1.64 | 0.69 | 2.39 | 0.02 | 0.69 | 0.54 | 1.28 | 0.20 |
|  | b1 | 0.17 | 0.29 | 0.58 | 0.56 | 0.19 | 0.27 | 0.71 | 0.48 | -3.01 | 1.58 | -1.91 | **0.06** | -0.17 | 0.80 | -0.22 | 0.83 | 0.74 | 0.58 | 1.28 | 0.20 |
|  | a2 | -0.18 | 0.33 | -0.55 | 0.58 | -0.15 | 0.30 | -0.50 | 0.62 | 3.10 | 1.81 | 1.71 | **0.09** | -0.17 | 0.82 | -0.21 | 0.83 | -1.19 | 0.84 | -1.42 | 0.16 |
|  | b2 | -0.24 | 0.35 | -0.68 | 0.50 | -0.25 | 0.32 | -0.76 | 0.45 | 1.96 | 1.92 | 1.02 | 0.31 | 0.09 | 0.91 | 0.10 | 0.92 | -0.10 | 0.85 | -0.12 | 0.91 |
| DTMAXabs | a1 | 1.43 | 0.15 | 9.43 | <0.01 | 1.55 | 0.14 | 10.77 | <0.01 | 1.20 | 0.59 | 2.04 | 0.04 | 2.10 | 0.49 | 4.32 | <0.01 | -0.60 | 1.05 | -0.58 | 0.56 |
|  | b1 | 0.02 | 0.14 | 0.16 | 0.87 | 0.09 | 0.13 | 0.65 | 0.52 | 0.55 | 0.65 | 0.84 | 0.40 | -0.45 | 0.40 | -1.12 | 0.26 | 0.11 | 0.14 | 0.75 | 0.45 |
|  | a2 | -0.05 | 0.22 | -0.23 | 0.82 | 0.05 | 0.20 | 0.23 | 0.82 | 1.28 | 1.22 | 1.05 | 0.29 | -0.69 | 0.60 | -1.14 | 0.25 | -3.21 | 1.92 | -1.68 | **0.09** |
|  | b2 | 0.05 | 0.21 | 0.22 | 0.83 | -0.02 | 0.19 | -0.11 | 0.91 | -1.51 | 1.31 | -1.16 | 0.25 | 0.47 | 0.52 | 0.90 | 0.37 | 1.09 | 0.55 | 1.98 | **0.05** |
| DTMOYabs | a1 | 1.41 | 0.18 | 7.89 | <0.01 | 1.53 | 0.17 | 9.04 | <0.01 | 0.88 | 0.64 | 1.39 | 0.17 | 2.10 | 0.54 | 3.89 | <0.01 | -0.15 | 0.88 | -0.17 | 0.87 |
|  | b1 | 0.03 | 0.14 | 0.21 | 0.84 | 0.07 | 0.13 | 0.58 | 0.56 | 1.58 | 0.92 | 1.71 | **0.09** | -0.32 | 0.37 | -0.86 | 0.39 | 0.08 | 0.24 | 0.36 | 0.72 |
|  | a2 | 0.27 | 0.40 | 0.67 | 0.50 | 0.25 | 0.37 | 0.68 | 0.50 | 0.49 | 2.67 | 0.19 | 0.85 | -0.07 | 1.02 | -0.07 | 0.94 | -5.78 | 2.26 | -2.55 | **0.01** |
|  | b2 | 0.43 | 0.47 | 0.91 | 0.36 | 0.22 | 0.43 | 0.52 | 0.60 | -3.49 | 3.87 | -0.90 | 0.37 | 1.13 | 1.08 | 1.04 | 0.30 | 4.49 | 1.62 | 2.76 | **<0.01** |
| MAXHum | a1 | 1.44 | 0.21 | 6.81 | <0.01 | 1.64 | 0.19 | 8.61 | <0.01 | 0.84 | 0.78 | 1.08 | 0.28 | 1.48 | 0.51 | 2.91 | <0.01 | 0.39 | 0.39 | 0.99 | 0.32 |
|  | b1 | -0.00 | 0.19 | -0.01 | 0.99 | -0.05 | 0.17 | -0.30 | 0.77 | 2.19 | 0.98 | 2.24 | **0.03** | 0.36 | 0.49 | 0.74 | 0.46 | -0.39 | 0.36 | -1.09 | 0.28 |
|  | a2 | -0.15 | 0.32 | -0.47 | 0.64 | -0.10 | 0.29 | -0.35 | 0.72 | 1.74 | 1.84 | 0.95 | 0.34 | -0.21 | 0.72 | -0.30 | 0.77 | -0.09 | 0.81 | -0.11 | 0.91 |
|  | b2 | -0.06 | 0.34 | -0.19 | 0.85 | 0.03 | 0.31 | 0.08 | 0.93 | -1.41 | 2.11 | -0.67 | 0.50 | -0.57 | 0.81 | -0.70 | 0.48 | 0.71 | 0.89 | 0.80 | 0.42 |
| MINHum | a1 | 1.42 | 0.22 | 6.34 | <0.01 | 1.60 | 0.20 | 7.86 | <0.01 | 0.68 | 0.79 | 0.86 | 0.39 | 1.58 | 0.56 | 2.81 | <0.01 | 0.25 | 0.40 | 0.62 | 0.53 |
|  | b1 | 0.02 | 0.19 | 0.12 | 0.91 | -0.01 | 0.18 | -0.05 | 0.96 | 2.43 | 1.07 | 2.27 | **0.02** | 0.22 | 0.52 | 0.43 | 0.66 | -0.16 | 0.37 | -0.42 | 0.68 |
|  | a2 | 0.21 | 0.43 | 0.49 | 0.62 | 0.12 | 0.40 | 0.30 | 0.76 | 3.07 | 2.55 | 1.21 | 0.23 | 0.50 | 1.07 | 0.47 | 0.64 | 1.61 | 1.23 | 1.31 | 0.19 |
|  | b2 | 0.38 | 0.52 | 0.73 | 0.47 | 0.23 | 0.47 | 0.50 | 0.62 | -0.38 | 2.72 | -0.14 | 0.89 | 0.67 | 1.22 | 0.55 | 0.58 | 2.31 | 1.36 | 1.70 | **0.09** |
| DIFFHum | a1 | 1.40 | 0.21 | 6.73 | <0.01 | 1.54 | 0.19 | 8.03 | <0.01 | 0.78 | 0.67 | 1.17 | 0.24 | 1.51 | 0.56 | 2.72 | <0.01 | -0.01 | 0.38 | -0.04 | 0.97 |
|  | b1 | -0.05 | 0.20 | -0.27 | 0.79 | -0.06 | 0.18 | -0.34 | 0.73 | -1.59 | 0.86 | -1.84 | **0.07** | 0.30 | 0.52 | 0.58 | 0.56 | -0.28 | 0.38 | -0.74 | 0.46 |
|  | a2 | 0.16 | 0.30 | 0.54 | 0.59 | 0.13 | 0.27 | 0.46 | 0.65 | 1.77 | 1.51 | 1.17 | 0.24 | 0.00 | 0.94 | 0.00 | 1.00 | 0.59 | 0.66 | 0.90 | 0.37 |
|  | b2 | -0.30 | 0.33 | -0.84 | 0.40 | -0.11 | 0.30 | -0.35 | 0.72 | 0.94 | 1.49 | 0.63 | 0.53 | -0.17 | 1.08 | -0.15 | 0.88 | -0.44 | 0.68 | -0.64 | 0.52 |
| MOYHum | a1 | 1.43 | 0.22 | 6.37 | <0.01 | 1.63 | 0.20 | 7.98 | <0.01 | 0.75 | 0.82 | 0.91 | 0.36 | 1.83 | 0.56 | 3.29 | <0.01 | 0.36 | 0.41 | 0.88 | 0.38 |
|  | b1 | 0.00 | 0.19 | 0.02 | 0.98 | -0.04 | 0.18 | -0.21 | 0.84 | 2.43 | 1.04 | 2.34 | **0.02** | 0.06 | 0.53 | 0.12 | 0.90 | -0.31 | 0.38 | -0.83 | 0.40 |
|  | a2 | -0.07 | 0.41 | -0.18 | 0.86 | -0.07 | 0.37 | -0.19 | 0.85 | 2.88 | 2.65 | 1.09 | 0.28 | 0.24 | 0.80 | 0.30 | 0.76 | 1.17 | 1.19 | 0.99 | 0.32 |
|  | b2 | 0.03 | 0.47 | 0.07 | 0.94 | 0.04 | 0.42 | 0.11 | 0.92 | -0.47 | 2.89 | -0.16 | 0.87 | -0.98 | 0.82 | -1.20 | 0.23 | 2.10 | 1.32 | 1.58 | 0.11 |

